# Molecular Dynamics Simulations of Bacterial Outer Membrane Lipid Extraction: Adequate Sampling?

**DOI:** 10.1101/2020.06.11.146001

**Authors:** Jonathan Shearer, Jan K. Marzinek, Peter J. Bond, Syma Khalid

## Abstract

The outer membrane of Gram-negative bacteria is almost exclusively composed of lipopolysaccharide in its outer leaflet, whereas the inner leaflet contains a mixture of phospholipids. Lipopolysaccharide diffuses at least an order of magnitude slower than phospholipids, which can cause issues for molecular dynamics simulations in terms of adequate sampling. Here we test a number of simulation protocols for their ability to achieve convergence with reasonable computational effort using the MARTINI coarse-grained force-field. This is tested in the context both of potential of mean force (PMF) calculations for lipid extraction from membranes, and of lateral mixing within the membrane phase. We find that decoupling the cations that cross-link the lipopolysaccharide headgroups from the extracted lipid during PMF calculations is the best approach to achieve convergence comparable to that for phospholipid extraction. We also show that lateral lipopolysaccharide mixing/sorting is very slow and not readily addressable even with Hamiltonian replica exchange. We discuss why more sorting may be unrealistic for the short (microseconds) timescales we simulate and provide an outlook for future studies of lipopolysaccharide-containing membranes.

## Introduction

In recent years, molecular models of bacterial membranes which incorporate much of the complexity and diversity of the lipids naturally found in them have become increasingly prevalent in molecular dynamics (MD) simulation studies. The relatively recent availability of appropriate atomistic/united atom parameters in commonly used force-fields, and more recently still, parameters for coarse-grained (CG) force-fields, have contributed to the rapidly increasing number of reported studies in which this lipidic diversity is considered^1–5^. Much of the attention to incorporating realistic lipid composition has focused on the complexity of the outer membrane (OM) of Gram-negative bacteria. These membranes are comprised of lipopolysaccharide (LPS) molecules in their outer leaflets and a mixture of phospholipids – (largely phosphatidyl-ethanolamine (PE), phosphatidyl-glycerol (PG) and cardiolipin (CDL) – in their inner leaflets. LPS molecules typically have between 4-6 acyl tails (depending on the species and adaptation state of the bacterium) and large headgroup regions comprised of multiple highly charged sugar moieties, which also vary between bacterial species. Here we focus on LPS from *E. coli*. Experimental and simulation studies have shown that LPS diffuses an order of magnitude slower than two-tailed phospholipids^6^, which for MD simulations of membranes containing LPS makes convergence problematic. For example, simulations of an atomistic *Pseudomonas aeruginosa* OM showed that equilibration of membrane properties only occurred after 500 ns^7^. Another study of protein-lipid interactions in *E. coli* OMs^8^, at the CG level or resolution, have shown that convergence is often observed within a single repeat, but not across multiple repeats. In some cases, even after 40 μs of simulation there was a lack of convergence between repeats. It was reasoned that strong inter-LPS interactions facilitated via cross-linking of divalent cations slows down the sampling of phase space during such simulations. Overcoming these sampling issues is of great importance in using MD as a predictive tool for the properties of bacterial envelope systems.

To understand the lipidic composition of a membrane it is of interest to consider the stability of a given lipid within a bilayer. This can be of particular importance when considering mechanistic processes of the membrane, such as defect formation or cell fusion. The stability of a lipid within a membrane can be quantified through the measurement of its free energy of extraction. As far as the authors are aware, lipid A is the only LPS lipid type for which the extraction free energy has been quantified to date^9,10^. This was performed in the context of immunological recognition of the ‘molecular pattern’ by mammalian Toll-like receptor 4 (TLR4), but in the context of membrane properties, lipid A is smaller than the minimal LPS molecule that is found to naturally occur in bacterial envelopes, namely the Re rough mutant of LPS (ReLPS), which includes a minimal core oligosaccharide.

Umbrella sampling (US) is an enhanced sampling method that has often been used to quantify the free energy profile or PMF along a reaction coordinate or collective variable (CV) of interest, such as e.g. membrane permeation^11,12^. One difficulty with the application of US to membrane-containing systems can be a lack of convergence, due to insufficient sampling times. Of particular interest was a previous atomistic study that focused on the reversible permeation of small solute molecules through a simple phospholipid membrane^13^. It was observed that during membrane permeation there were slow modes of reorganization that occurred over hundreds of nanoseconds. However, such timescales often prove too computationally expensive, and so in the present study, the molecular models were simulated at the CG level using the popular Martini force field^14–16^. The validity of CG models to study relative free energy landscapes of protein-lipid interactions has been proven by multiple studies^17–19^.

In the present study, we apply US to calculate PMFs for the extraction of lipids from both Gram-negative bacterial OM and inner membrane (IM) models. The convergence of the resultant PMFs tend to be deficient for the extraction of LPS molecules, in contrast with phospholipids, due to the sampling of variable extraction paths resulting from poor lipid sorting. We demonstrate that this may be alleviated via the decoupling of lipid extraction from divalent cations which cross-link the LPS headgroups. Furthermore, we assess whether the problematic issue of slow lipid sorting might be resolved in future using enhanced sampling protocols based on Hamiltonian replica exchange (HREX) MD.

## Results

In the following sections, we outline the application of US to calculate PMFs for extraction of lipids from both Gram-negative bacterial OM and IM models, using a range of CVs (Table 1). The OM model was comprised of an extracellular leaflet of ReLPS and an inner leaflet of 90% 16:0–18:1 phosphatidyl-ethanolamine (POPE), 5% 16:0–18:1 phosphatidyl-glycerol (POPG) and 5% CDL, with LPS lipids neutralized by calcium ions. In comparison, the IM composition was 75% POPE, 20% POPG and 5% CDL. Excess charge due to phospholipids in either system were balanced out with sodium ions. An initial production run of up to 10 μs was run for each membrane model to improve the convergence of membrane properties. Further details on the simulation models are given in the Methods section.

**Table 1.** Summary of all US calculations carried out for OM and IM systems. For all OM systems, 3 different ReLPS molecules were independently extracted, whilst the extraction of one of those lipids was performed in triplicate (thus yielding a total of 5 PMFs). For all IM systems, 3 different lipid molecules were independently extracted.

| Membrane system | PMFs calculated | CV | US time per window (μs) |
| --- | --- | --- | --- |
| OM | 5 x ReLPS | COM_MEMB | 10.0 |
| OM | 5 x ReLPS | COM_CY | 7.0 |
| OM | 5 x ReLPS | HEAD_MEMB | 7.0 |
| OM | 5 x ReLPS | HEAD_CY | 7.0 |
| OM | 5 x ReLPS | COM_CY_deCA | 3.0 |
| OM | 3 x POPE | HEAD_CY | 1.5 |
| OM | 3 x POPG | HEAD_CY | 2.0 |
| OM | 3 x CDL | HEAD_CY | 2.0 |
| IM | 3 x POPE | HEAD_CY | 1.5 |
| IM | 3 x POPG | HEAD_CY | 2.0 |
| IM | 3 x CDL | HEAD_CY | 2.0 |

### Umbrella sampling: the extraction of lipopolysaccharides

A single ReLPS was extracted from one leaflet using steered MD (SMD), in order to carry out US and hence calculate its PMF (Table 1). This process was independently repeated in triplicate for the same ReLPS molecule, in order to assess convergence. Previous work for such calculations has shown convergence of physical properties within a single repeat, but often not across different repeats^8,20^. Thus, two additional lipids were also randomly selected from the same leaflet for extraction and for PMF calculations; it was reasoned that the PMF profiles of all lipids of the same type should be the same, in the limit of complete convergence of the US simulations. Each of the PMFs for ReLPS extraction were calculated using four different trial CVs, in order to determine their role in PMF convergence. In each case, the CV was based on the z-component of the distance between the center of mass of selected reference groups for the extracted lipid and for parts of the remainder of the membrane. The reference group of the extracted ReLPS molecule was either its entire center of mass (‘COM’) or the center of mass of the phosphate beads (‘HEAD’). The reference group for the membrane was either its entire center of mass (‘MEMB’) or the center of mass of all lipid beads within a cylindrical projection around the extracted lipid (‘CY’). Thus, the CVs in order of the cumulative size of each reference group were: ‘COM_MEMB’, ‘HEAD_MEMB’, ‘COM_CY’ and ‘HEAD_CY’. Each PMF was calculated based on up to 10 μs of sampling per window (Table 1).

The first step in assessing the efficiency of an US simulation was to measure the coverage of the reaction coordinate by the umbrella windows. To this end, the percentage overlap between adjacent umbrella windows was measured for the extraction of ReLPS with each CV (Figure S1). Generally, the overlap between adjacent windows was between ~10-70% across PMFs, with an average of around 35%. A previous US study of the poration of ions through ion channels determined that a 5% overlap between adjacent windows provided sufficient coverage of the reaction coordinate^21^. Each set of percentage overlaps was compared to the ideal overlap between two harmonic biases spaced 0.1 nm apart (34%). The ideal overlap was determined using Equation 1 (see Methods section).

Now the convergence between different ReLPS extractions will be discussed. For each CV, three different ReLPS lipid molecules were extracted, with one of them (‘lipid 1’) being extracted in triplicate US calculations. Thus, convergence could be probed for repeated PMFs of a single lipid versus different lipids. All PMFs were calculated across the last 2 μs of each umbrella window (Figure 1). The ReLPS extraction free energies were spread across a large range of 35-55 kcalmol^-1^. There was little convergence evident when comparing between the PMFs of repeat extractions or different ReLPS lipids. The COM_MEMB CV had 10 μs of umbrella sampling run for each window, but only two out of three extractions of the same lipid were similar. The CV with least similarity between different PMFs was HEAD_MEMB. The only CV that gave the same PMF profile for all three repeats of a single lipid extraction (lipid 1) was COM_CY. Furthermore, two out of the three ReLPS lipids extracted with the COM_CY CV gave the same lipid extraction free energy.

**Figure 1.**
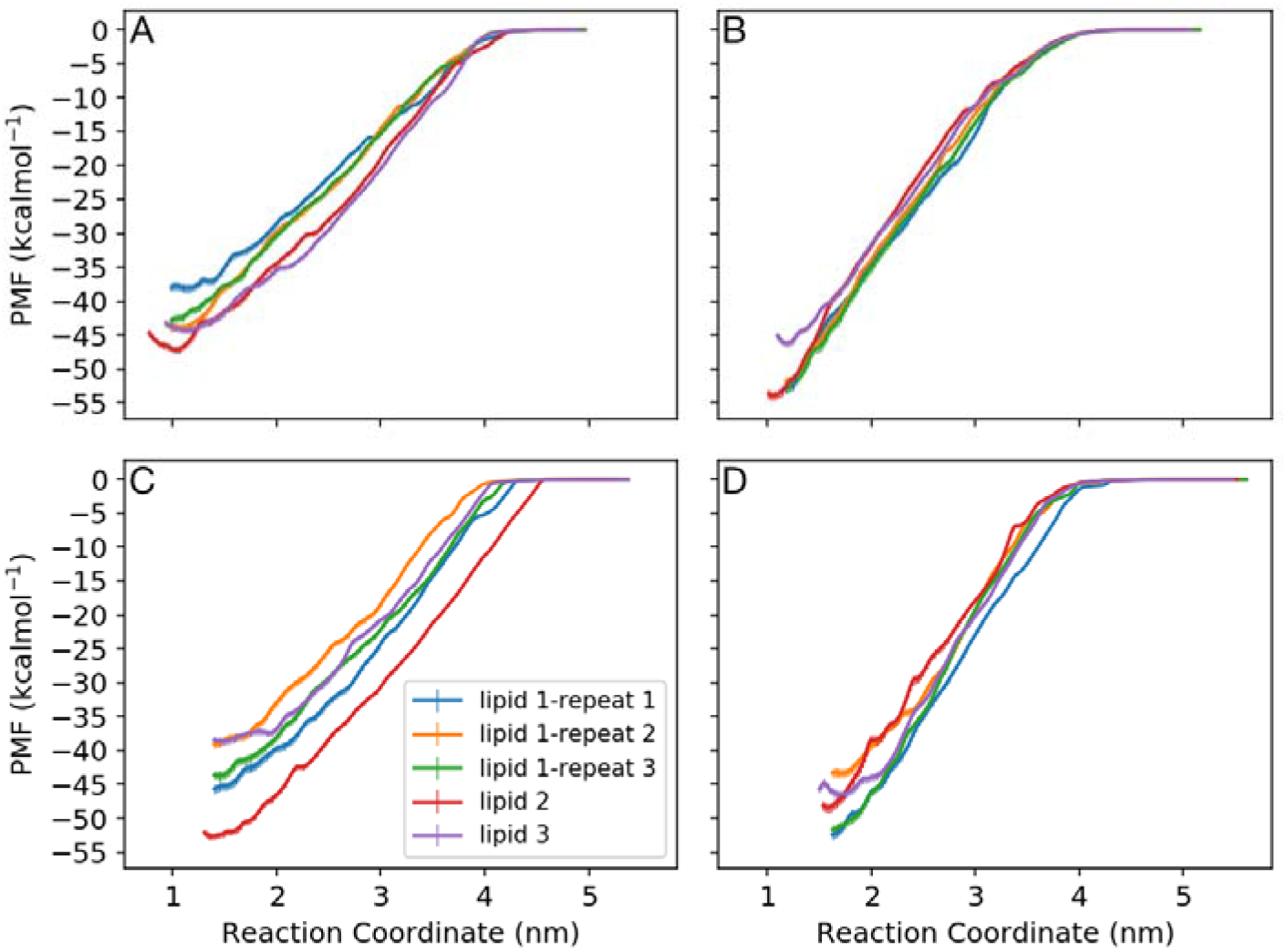
PMFs of ReLPS extraction from the OM using different CVs. The CVs included A) COM_MEMB, B) COM_CY, C) HEAD_MEMB and D) HEAD_CY, measured across the last 2 μs of each window. The error bars shown were calculated with WHAM using a bootstrapping procedure.

One point of interest is whether it may be shown that individual PMF profiles are ‘internally’ converged, without comparison to any other PMFs. The convergence of a single PMF was assessed by calculating its profile across a series of adjacent time interval blocks; in this case, PMFs across 1 μs intervals were measured every 0.5 μs. If convergence is reached, then these profiles should begin to oscillate around an average or tend towards a fixed profile shape. Such internal convergence could typically be reached for all lipid extraction calculations, as exemplified by the COM_CY CV system (Figure 2).

**Figure 2.**
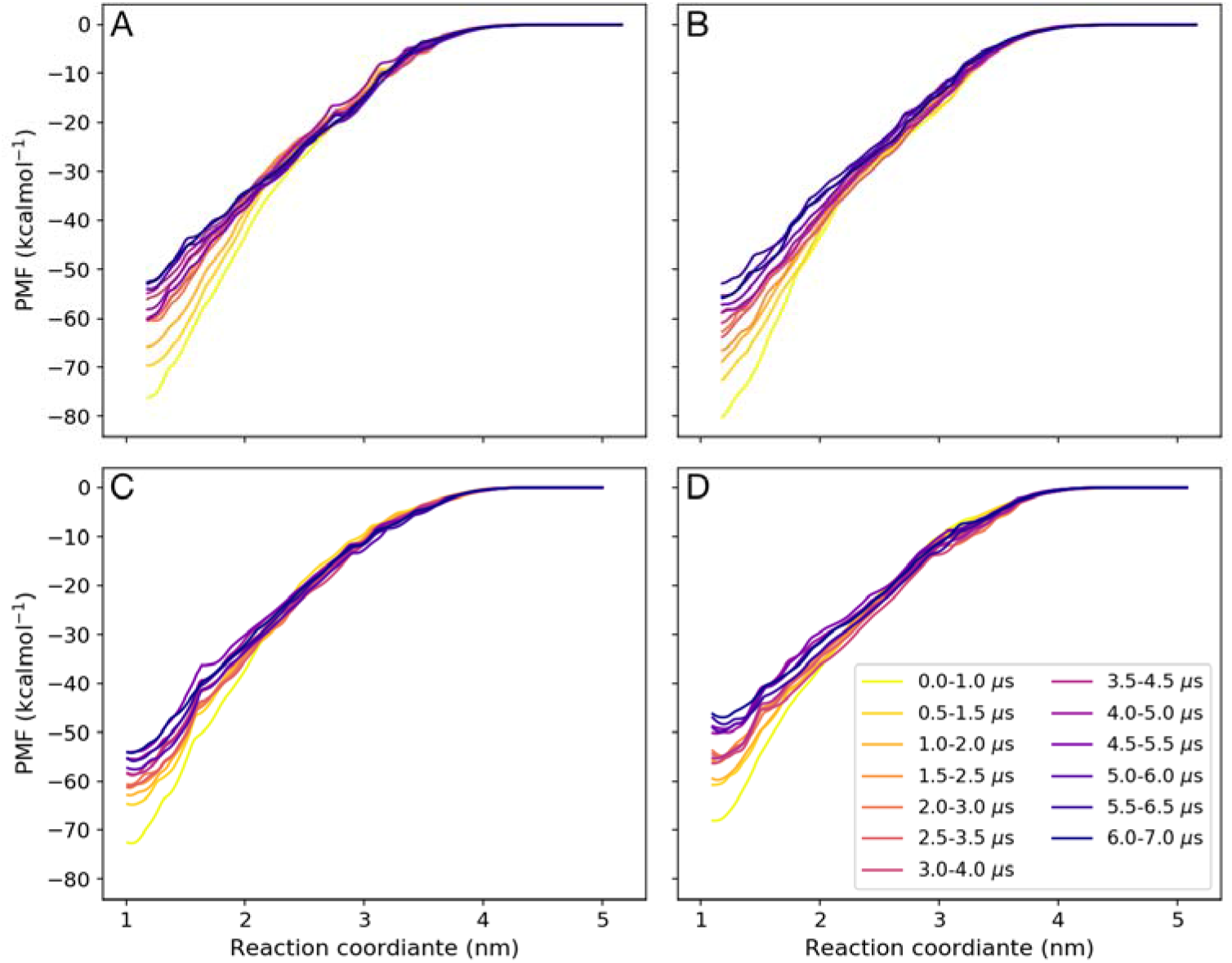
PMF convergence. PMFs of ReLPS extraction for A) lipid 1-repeat 1, B) lipid 1-repeat 2, C) lipid 2 and D) lipid 3 (see Figure 1) using the COM_CY, measured across 1 μs blocks.

Bootstrap methods that incorporate autocorrelation times can underestimate the uncertainty, especially if sampling is limited. To see if the trends discussed here were consistent across free energy calculation methods, the PMFs across the last 2 μs of US simulations were recalculated using the multistate Bennett acceptance ratio^22^ (MBAR) method (Figure S2). Overall there was little change in the observed trends when using different methods, but the estimated uncertainties were slightly larger (though still only on the order of 1 kcal mol^-1^ or less).

### Umbrella sampling: membrane reorganization

It is clear that in many cases the ReLPS extraction free energies do not converge – when comparing PMFs for replicas of the same lipid molecule, different lipid molecules, or different CVs – but what is not clear is why. It is thus of interest to investigate the differences between membrane reorganization and the path of the extracted lipid. The integrated autocorrelation time (IACT) of each window was measured for each repeat across all CVs (Figure S3). The IACT was very inhomogeneous across each reaction coordinate, ranging from a few nanoseconds to over 800 ns. In particular, the IACT was always larger towards the core of the lipid bilayer, an observation which mirrors the large barriers to sampling in the membrane previously observed for DOPC bilayers^13^. measurement of the IACT confirmed that the PMFs in LPS systems even in CG representation should be calculated over microsecond times scales to ensure that a significant number of CV values are uncorrelated. In previous work, we showed that the on/off rates of LPS from the surface of OMPs stretch beyond tens of microseconds^8^, and this lack of LPS sorting correlates with the difficulties observed here in attaining PMF convergence between different ReLPS extraction events.

We attempted to assess the performance of each CV in assisting good sampling of membrane properties during PMF calculation. First, the difference in path sampling across CVs was explored. There was one big difference between the lipid extraction paths for CVs that used the COM and HEAD reference groups. When the COM reference group was used, the lipid had more rotational and translational degrees of freedom for each CV value, compared to CVs that included the HEAD reference group. When a lipid was extracted using the COM group, the lipid would rotate such that the LPS sugars and lipid A head group were in contact with the membrane for as long as possible (Figure 3). Use of the HEAD reference group pulled the lipid out of the membrane without allowing this flip to easily occur.

**Figure 3.**
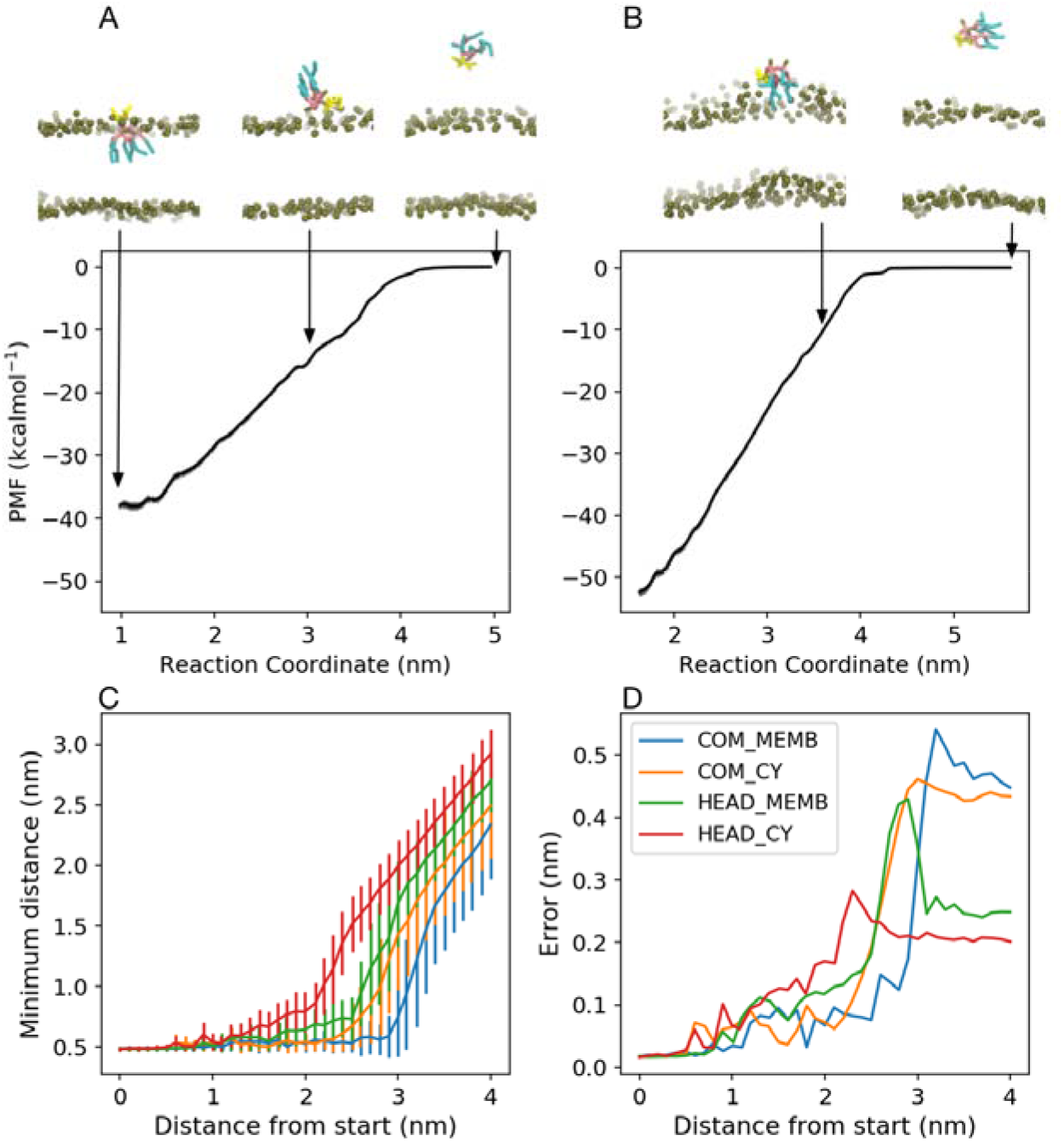
Extraction of ReLPS from the OM. Snapshots along the pathways of extraction are shown above their corresponding PMF for A) COM_MEMB and B) HEAD_CY CVs. Key for snapshots: gold=phosphates, yellow=sugars, pink=acyl groups, cyan=lipid tails. C) Minimum distance from the phosphate beads in the extracted ReLPS to the OM, and D) standard deviation in the minimum distance relative to the start of each CV.

The minimum distance between the phosphate beads of the extracted LPS and the membrane was calculated to show the path that the lipid headgroup took for each CV (Figure 3C). This confirms that lipid A left the membrane surface much later in the reaction coordinate when the COM reference group was used. As the ‘specificity’ of the CV increased, there was a decrease in the standard deviation of the minimum distance when ReLPS exited the membrane (Figure 3D). For the HEAD_CY CV there were three distinctive regions: (i) a flat plateau when lipid pulling began (<1 nm from the initial, equilibrium position); (ii) a gradual increase in the minimum distance as the lipid headgroup was extracted from the membrane (<2 nm); and (iii) a final plateau when the entire lipid was in solution (<2.5 nm). These three regions were only observed when the lipid was pulled from the phosphate beads. What these results show is that the COM reference group imparts the lipid with more flexibility during the extraction process. Thus, CVs that utilized the COM reference group should result in longer convergence times, but may capture the minimum free energy path of lipid extraction more accurately.

To further assess the reorganization of the membrane across repeats, the orientation of the lipid during the extraction process was considered. This was quantified by measuring the angle of the core sugars and lipid A headgroup of each ReLPS molecule with respect to the z axis (*θ*_LPS_), and the deviation order parameters within a 1 nm radius (in the xy plane) of the extracted lipid from that of a neat membrane 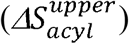, each as a function of reaction coordinate (Figure 4). This analysis was only performed for the COM_CY and HEAD_CY CVs to compare and contrast the lipid path and local membrane reorganization when convergence, or a lack thereof, was obvious. During most lipid extractions, there was a small perturbation in the local lipid tail order. The distribution of 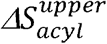 values for repeat extraction of lipid 1 using the COM_CY CV were similar, but dissimilar to the extraction of lipid 3 (Figure 4A and C). Therefore, the trends in local membrane reorganization seemed to mirror the convergence of PMFs shown in Figure 1. Similarly, the *θ_LPS_* values for the repeat extractions of lipid 1 were more similar compared to the extraction of lipid 3. One noticeable difference was that during the initial part of the extraction of lipid 3 (the first 1.5 nm) the LPS headgroup tilted such that it lay parallel to the membrane normal. Even larger differences were noted when comparing between the HEAD_CY and COM_CY CVs. The variation of the ReLPS headgroup orientation was greatly reduced for the HEAD_CY CV. When the HEAD_CY CV was used, the orientation of the headgroup changed in a linear manner from 45-150°, until the ReLPS lipid broke free from the membrane surface and could freely rotate in solution. This linear increase in orientation was consistent with the lipid headgroup flipping to maximize contacts with sugar groups during the extraction process. In this way, the lipid is pulled out headgroup first, and as the tail is pulled out, the headgroup flips to contact the sugars. In summary, the variable extraction paths associated with different CVs are associated with poor convergence of PMFs between systems due to slow LPS lateral sorting.

**Figure 4.**
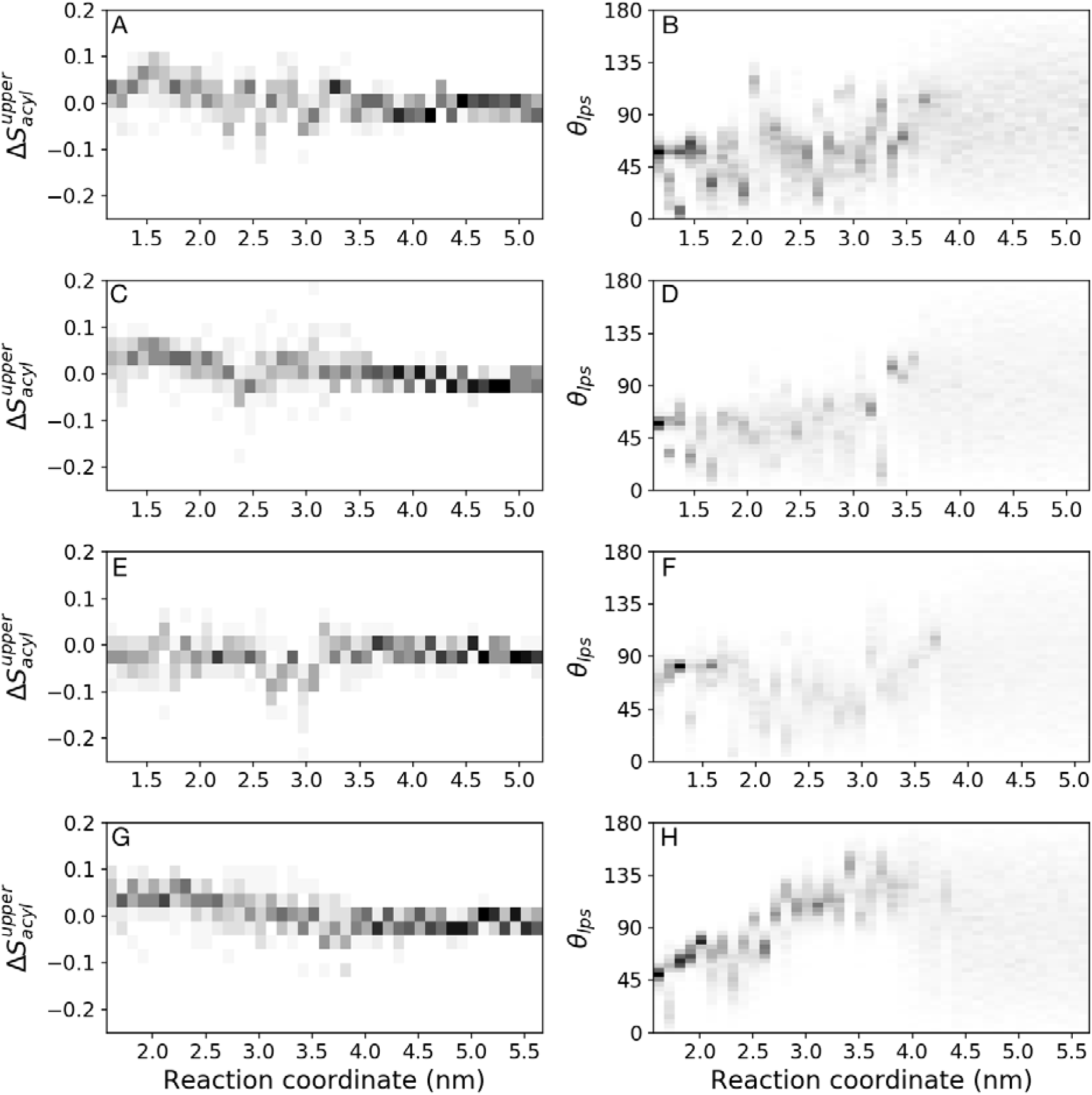
Orientation of LPS molecules during extraction. (left) Mean acyl order parameters for LPS within 1 nm of the extracted lipid in the xy plane, compared to that in an unbiased bilayer, 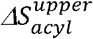. (right) Orientation of the LPS headgroup compared to the bilayer normal, *θ_LPS_*. Analysis was carried out for the extraction of ReLPS using the COM_CY CV for (A and B) lipid 1 – repeat 1, (C and D) lipid 1 - repeat 3, (E and F) lipid 3 and the HEAD_CY for (G and H) lipid 1 – repeat 1. Each analysis was carried out for the entire umbrella sampling trajectory (Table 1).

### Umbrella sampling: extracting phospholipids

For the sake of comparison, phospholipids in the lower leaflet of the OM and IM were extracted using the HEAD_CY CV (Table 1), where the reference group of the lipid was the center of mass of the phosphate beads, or the glycerol linker in the case of CDL. For each membrane, three different lipids were extracted independently to yield separate PMFs. In all cases, it was observed that convergence in PMFs between different lipids of the same type was achieved more quickly than the LPS systems, within 2 μs (Figure S4). The difference in the free energy of extraction between the inner leaflet of the OM and the IM was −0.35, 1.24 and 1.70 kcal mol^-1^ for POPE, POPG and CDL, respectively (standard deviations < 0.2 kcal mol ^-1^). It is interesting that only the stabilities of POPG and CDL were increased by the presence of ReLPS. The increased stability of CDL in the membrane may be explained by the previously observed clustering of CDL below areas of low ReLPS sugar density^20^.

### Umbrella sampling: decoupling ion ‘cross-links’ from LPS extraction

A key difference between the leaflets containing phospholipid versus LPS lies in the structural role of ions. Both computational and experimental studies have shown that the headgroups of LPS lipids form strong ionic interactions via divalent cations^6,23,24^, which restricts lipid diffusion. We therefore reasoned that the coupling of divalent cations between the ReLPS molecule being extracted and the remainder of the membrane could be a source of the observed variation in extraction paths across separate calculations. This would help to rationalize the observed range of variable ReLPS free energy profiles, in contrast with the rapidly converging phospholipid PMFs. To assess this, we first performed a series of SMD simulations for the COM_CY CV using a range of velocities (1, 0.1, 0.05, or 0.01 nm ns^-1^), with the ReLPS molecule reaching a distance of 4 nm from the membrane over a timescale of 4, 40, 80, or 400 ns, respectively. In all these simulations, even at the slowest pulling speed, divalent calcium ions were found to remain electrostatically bound to the anionic constituents of ReLPS upon their complete extraction into solvent (Figure 5A, B). Therefore, we next ran SMD simulations but with all membrane-bound calcium ions restrained in the z-dimension to prevent their coupled extraction. We subsequently used the resultant coordinates of the extracted calcium-free ReLPS molecules to perform umbrella sampling calculations (‘COM_CY_deCA’, Table 1), again for one lipid molecule in triplicate and once for two other lipid molecules. Based on block analysis, as shown in Figure 5C-F, the PMFs for all systems individually converged within 3 μs of sampling per window. Strikingly, the PMFs for replicas for the same lipid or for separate lipids all yielded a final free energy of extraction of −50 to −55 kcal mol^-1^. Thus, even when using the COM reference group, which we have already shown may result in longer convergence times, decoupling of the cross-linking divalent cations from LPS extraction significantly helps to improve PMF convergence properties.

**Figure 5.**
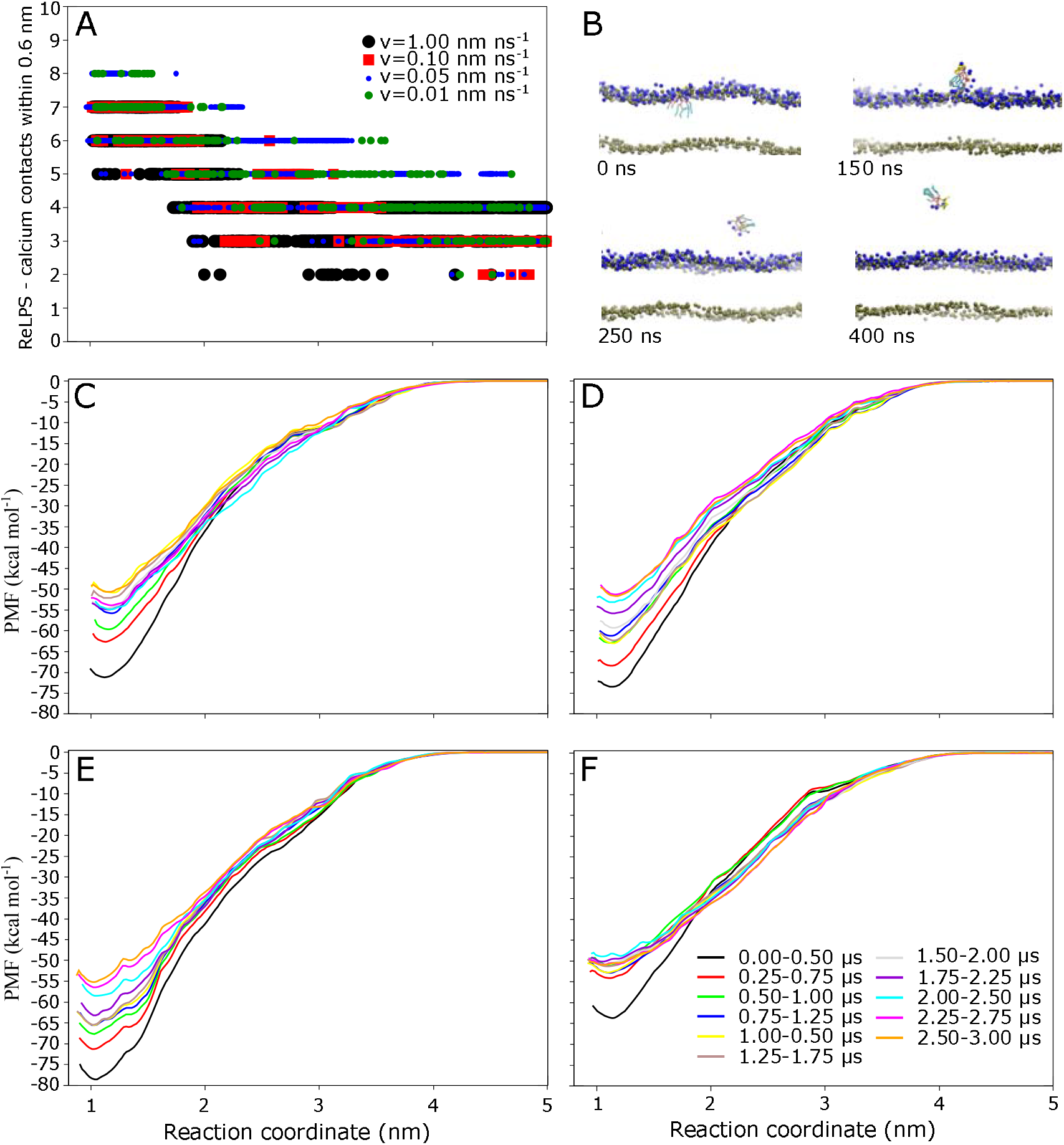
Role of calcium in SMD and PMFs for ReLPS extraction using the COM_CY CV. A) Contacts within 0.6 nm between ReLPS and calcium ions along the reaction coordinate during SMD simulations with ranging velocities. B) Snapshots during SMD at pulling rate of 0.01 nm ns^-1^, highlighting calcium coordination of extracted lipid. Key for snapshots: gold=phosphates, yellow=sugars, pink=acyl groups, cyan=lipid tails, blue=calcium ions. PMFs are shown for C) lipid 1-repeat 1, D) lipid 1 - repeat 2, E) lipid 2, F) lipid 3, measured across 0.5 μs blocks of umbrella sampling.

### Enhanced lipopolysaccharide mixing via HREX MD

As shown above, in the absence of artificial constraints to decouple divalent cations from ReLPS in order to simplify the extraction path, incomplete PMF convergence results, at least in part, from the lack of LPS lateral sorting. Thus, we next tested the capacity for HREX MD to enhance the lateral mixing of ReLPS in the OM model. In HREX, several replicas of the same system are simulated whilst a specific portion of the Hamiltonian is tempered according to λ, and coordinates are exchanged from time to time with a Monte Carlo procedure. When designing a HREX protocol, it is important to first carefully consider what should be the tempered region. A variety of trial tempering regions of the ReLPS molecules were therefore first tested in 4 μs HREX simulations (Figure S5), during which no replicas were exchanged (see Supplementary Methods for further details). Nearest neighbor analysis was performed to quantify lateral sorting of LPS as a function of λ. For LPS to be well mixed, two conditions were assumed to be necessary: (i) an LPS molecule must be a nearest neighbor with every LPS in the membrane during the course of a trajectory; and (ii) an LPS molecule must have an equal probability of being neighbors with any other LPS in the same leaflet. To quantify these two assumptions, two metrics were designed, the NN index and *RMSD_NN_*. The NN index was a value between 0 and 1 that quantified the first assumption. If the NN index was 0 then the neighbors of a single LPS never changed. Conversely, if the NN index was 1 then the LPS had been neighbors with every other LPS in the membrane across the analyzed trajectory. The *RMSD_NN_* was defined as the RMSD of nearest neighbor frequencies for a single LPS, and would be 0 if the second assumption for ideal mixing was satisfied. These two metrics were used to determine that the ideal tempering region was the lipid A headgroup plus sugar moiety components of the ReLPS molecules, in which the mixing was only slightly less than when using the entire lipid (Figures S5, S6). The group may help to improve lateral mixing by reducing the tendency for Martini carbohydrate units to aggregate^25,26^. For subsequent HREX studies, this tempering region was thus used, with the λ range set to 1.000-0.645, the number of replicas set to 24, and the exchange attempt interval (EAI) set to 0.5 ns.

To assess the lateral sorting of ReLPS in HREX simulations, the NN index and *RMSD_NN_* were measured across time intervals of increasing lengths (Figure 6). After 25 μs, the ReLPS lipids in the ground state replica had an average NN index of ~0.65. When λ < 0.85 then every LPS lipid had been neighbors with every other LPS lipid. While *RMSD_NN_* tended to 0 at lower λ values, mixing was quite inhomogeneous at higher λ values. As the simulation time progressed, there were diminishing returns in the increase of ReLPS sorting and thus it is unclear how much additional simulation time would be required for homogenous mixing of the ground state.

**Figure 6.**
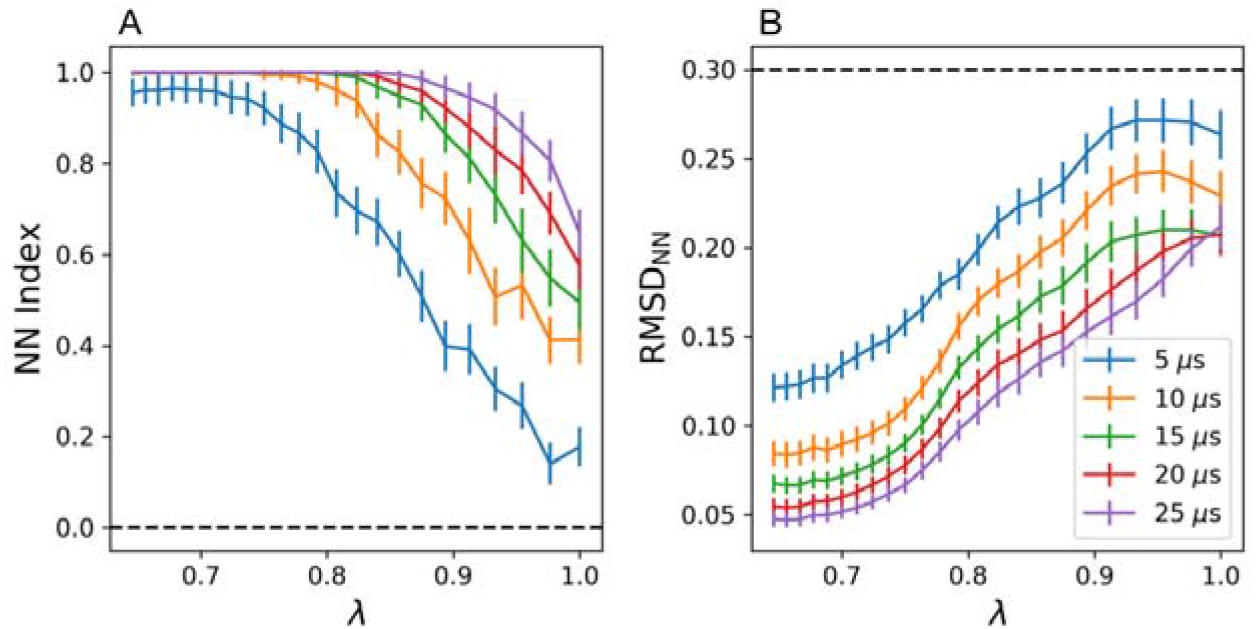
Replica mixing in HREX simulations. A) Nearest neighbor index (NN Index) and B) *RMSD_NN_* versus λ for ReLPS in an OM model (8×8 nm in the xy plane) across different time intervals. The dotted lines represent the values obtained when no LPS mixing occurred.

To measure the convergence of membrane properties in different areas of replica space, the area per lipid (APL) of ReLPS and the membrane thickness of the two replicas at the extremes of λ space was calculated (Figure 7). The APL and membrane thickness had converged for the ‘hottest’ replica, but with large errors in the block averages. The errors for the membrane properties of the maximum replica were larger than those of the ground state replica. This is consistent with faster kinetics and LPS mixing for the maximum replica. The membrane properties of the ground state replica converged after 14 μs of simulation.

**Figure 7.**
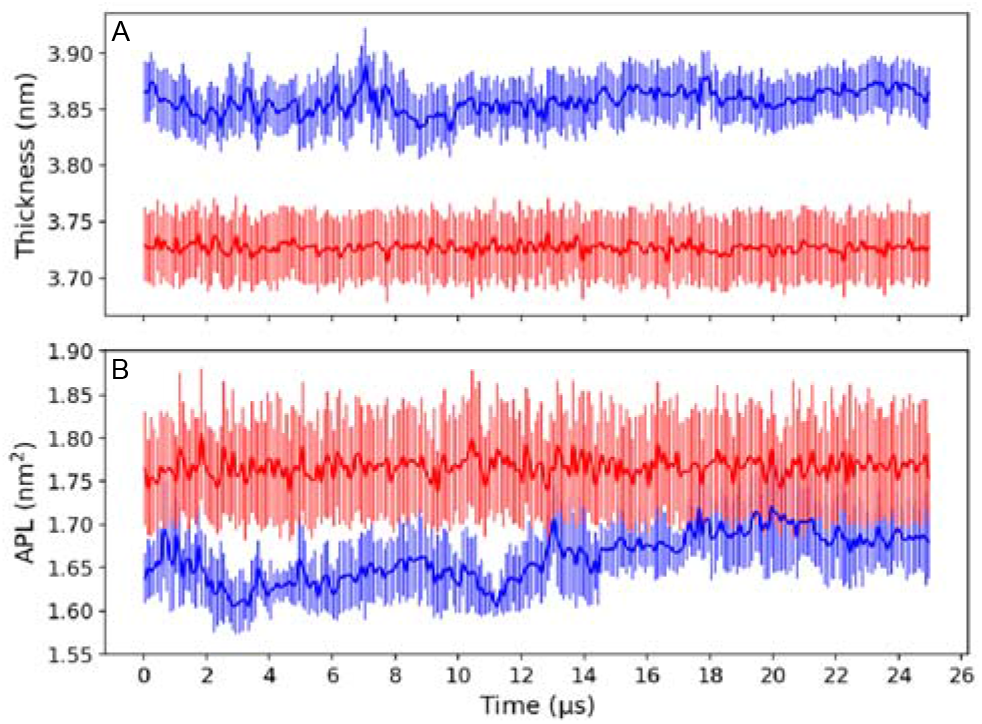
Membrane properties during HREX simulations. A) Membrane thickness and B) APL of ReLPS vs time for the maximum (red) and minimum (blue) replicas. Averages and errors determined from the block average and standard deviation, respectively, across 100 ns intervals. The APL and membrane thickness were calculated using the center of geometry of any phosphate beads in each lipid with the FATSLiM program ^27^.

To further probe the convergence of lipid lateral sorting, the Block Covariance Overlap Analysis Method (BCOM) method was used to measure the covariance overlap between the PCA modes in different blocks of a trajectory, and to quantify the relaxation speed of the system^28^, based on the positions of ReLPS phosphates (see Supplementary Methods for further details). For all block sizes, this analysis confirmed that HREX MD significantly improved the relaxation of the system (Figure S7). However, complete convergence was not achieved even for the largest block size, and it is difficult to estimate the amount of additional simulation time required, given the aforementioned diminishing returns in sampling improvements with respect to time.

## Discussion

The results of the present study highlight important points to consider when designing future MD studies of bacterial envelope models, particularly regarding the slow rate of convergence of simulation studies involving LPS. Here umbrella sampling was using to extract LPS from the bacterial OM, and phospholipids from the OM inner leaflet or IM. The free energy of extraction of ReLPS was determined using up to 10 μs of simulations per umbrella sampling window. There was minimal convergence between the free energy of extraction of a single ReLPS or different lipids across a range of CVs. In fact, there was only one CV for which all three repeats of the extraction of a single LPS resulted in the same extraction free energy. In a few cases the extraction of different LPS lipids converged to unique extraction free energies. Furthermore, the most specific CV was not the CV that performed the best. It seems feasible that further optimization of the CV may improve convergence for repeat extractions of the same LPS molecule. In particular, the further development of automated CV optimization with machine learning techniques^29^ is of key importance for future free energy studies of complex systems.

Subsequent US calculations focused on the OM inner leaflet or IM revealed that convergence could be achieved, and in a comparatively short timeframe compared to the LPS leaflet. It was also found that the extraction of PG and cardiolipin from the OM was more energetically demanding than the IM; this may be related to the LPS induced clustering of cardiolipin in the lower leaflet of the OM, as observed in a previous study^20^, and the overall anionic nature of the headgroups (compared to the zwitterionic PE). Irrespective, we noted that a key difference between phospholipid-versus LPS-containing leaflets is the role of ‘structural’ divalent cations. Further investigation confirmed that decoupling of calcium from the ReLPS molecule being extracted helped to achieve more reproducible reaction paths between separate PMFs, as well as speeding convergence rates similar to those for the phospholipid calculations.

In the absence of artificial constraints to decouple divalent cations from ReLPS during the initial extraction process, the lack of convergence for the associated PMFs likely result from the restricted lateral sorting and mobility of lipids. The lateral sorting of phospholipids was much faster, which was reflected by the convergence of their PMFs within 2 μs. In an attempt to improve lateral sorting, HREX MD was applied to ReLPS in an OM model, generating 25 μs of simulation per replica. This improved lipid sorting and system relaxation over the equivalent unbiased simulation. In this regard, the application of HREX MD to the OM was successful. However, the long autocorrelation times of LPS-containing systems resulted in replica mixing that is extremely inefficient and occurs over microseconds. It has been noted in the literature that the EAI should be larger than the IACT of the potential energy of the tempered region^30,31^, or inefficient sampling (‘exchange trapping’) may result. Unfortunately, based on additional unbiased 2 μs production runs, we estimate the IACT to be ~900 ns, which would be a computationally unfeasible EAI, and an EAI of 0.5 ns was thus utilized during each HREX simulation. To avoid trapping in certain areas of replica space, the exchange probabilities for each replica trajectory should not vary greatly^30^. Analysis of replica mixing of our trajectories (see Supplementary Methods for further details) revealed exchange probabilities comparable to those of previous studies^32^ (Figure S8A), but the total residency time varied greatly across replica space, with exchange trapping often occurring for a few replicas at the extremes of replica space (Figure S8B). Replica mixing (Figure S8C) and global round trips per replica (Figure S8D) were greatly reduced compared to other HREX studies^32,33,30,33,34^, and using previous replica simulations^30^ as a benchmark, we estimate that hundreds of microseconds of simulation would be required for sufficient mixing to be achieved, which is unfeasible for all but the smallest of systems due to computational requirements – and frustratingly, it is the much larger systems that are of biological interest. Further optimization of the HREX parameters could be investigated, but given the long IACTs of LPS, it is possible that HREX may prove unsuitable for such systems.

Another question, that has not been addressed here, is how much lateral sorting should LPS undergo? Some experimental data for smooth LPS suggests LPS domains can persist for days in phospholipid membranes^35^. Furthermore, it seems that LPS molecules are tightly coupled to OMPs^36,37^. In addition, currently there is limited experimental data available for the diffusive behavior of LPS (although increasingly experimentalists are beginning to address this, and thus in future, data will become available) and consequently one should consider whether HREX MD may result in unphysical levels of mixing. Thus, we are left with the conclusion that the LPS mobility we are achieving with our CG simulations is accurate for the timescales we are probing and thus short timescale phenomena will be as representative in these membranes as in any other, but the simulation timescales that we can currently achieve in practice are not long enough for convergence of free energies. For example, the extent of *change* in the conformations of the loops of OMPs within a certain timeframe will be represented well. In contrast, calculation of the free energy of interaction of those loops with LPS will take a lot more simulation to converge. Consequently, the design of future simulation studies of LPS-containing membranes requires some thought, and it is highly likely that there will not be a single ‘one size fits all’ approach, but rather the models and methods will depend on the questions being asked. One might envisage reparametrized CG models in which the sugar moieties of LPS undergo (unrealistically) faster conformational rearrangements – simply as a way of converging the free energies of association with proteins (the kinetics would be unreliable). Other methods to study, for example, the movement of protein-LPS complexes may include addition of phospholipids in the outer leaflet to ‘loosen’ the LPS-LPS interactions, or perhaps to reduce counterion-phosphate interactions via e.g. scaling down of charges, or e.g. their ‘uncoupling’ using restraints, as demonstrated in the present study. These approaches will of course have their own limitations and are included here by way of encouraging the reader to think about future design and composition of simulations of bacterial membranes to appropriately address the scientific questions.

## Experimental Procedures

### Membrane models

The OM model consisted of an upper leaflet of ReLPS and a lower leaflet of 90% POPE, 5% POPG and 5% CDL. ReLPS refers to the Re rough mutant of LPS, which is comprised of a lipid A moiety and a core oligosaccharide component containing two KdO units^38^. The second membrane studied was the symmetric IM (75% POPE, 20% POPG and 5% CDL). The ReLPS model was parameterized in previous work by the Khalid group^38^. The initial coordinates of each system were generated with the CHARMM-GUI web interface^39,40^. LPS lipids were neutralized with calcium ions and then any remaining charge was balanced out with sodium ions.

### Simulation Parameters

All simulations were run with the GROMACS MD package^41^ (version 2018) and the Martini (version 2.2) force field^14–16^. HREX was run by patching GROMACS with plumed (version 2.4)^42,43^. Simulations were run at a temperature of 323 K and a pressure of 1 bar. The temperature was controlled with the stochastic velocity rescale thermostat^44^ with a coupling constant of 1 ps. Equilibration steps used the Berendsen^45^ semi-isotropic barostat with a coupling constant of 4.0 ps, whereas for production runs the Parrinello-Rahman^46^ barostat with a coupling constant of 12.0 ps was used. A timestep of 10 fs was used for all enhanced sampling simulations; in all other cases the timestep was 20 fs. The short range cutoff for non-bonded interactions was 1.2 nm and utilized the Potential-Shift-Verlet cutoff scheme. The electrostatic interactions were determined with reaction field with short and long range dielectric constants of 15 and 0 (infinite shielding), respectively.

### Simulation setup: Umbrella sampling

The OM and IM systems used in the US simulations were generated with CHARMM-GUI (dimensions 12×12 nm^2^ in the xy plane and 4 nm water above and below the membrane). Each system was then relaxed with a series of minimizations followed by up to 30 ns of equilibration, following which additional solvent was added to give a box height of 20 nm. Following minimization and equilibration, a 10 μs or 5 μs production run was carried out for OM and IM systems, respectively. Once the production runs of the unbiased simulations were completed, a number of lipids were randomly selected for extraction. It is worth noting that the same lipids were extracted across all CVs. For each CV, a single ReLPS extraction was repeated 3 times, to give a total of 5 ReLPS extractions per CV. SMD was used to pull a lipid out of the membrane, with a harmonic bias of 1000 kJmol^-1^ nm^-2^ and a pull rate of 0.5 nm ns^-1^ for each CV. Additional SMD simulations for the COM_CY system were also extracted with a range of other velocities (1, 01, 0.05 and 0.01 nm ns^-1^) to test the role of calcium cross-linking upon the extraction pathway. Next, SMD simulations were run with a pulling velocity of 0.02 nm ns^-1^ whilst applying position restraints to calcium ions in the xy plane (force constant of 200 kJ mol^-1^ nm^-2^). Then, for all systems, the initial coordinates of 41 windows were extracted every 0.1 nm along the SMD path, for subsequent US. For each US window, a harmonic bias of 1000 kJmol^-1^ nm^-2^ was applied to restrain the lipid around the initial CV value, and a production run of up to 10 μs was generated per window (see Table 1).

All PMFs were calculated with the WHAM algorithm implemented within GROMACS^47^ (unless state otherwise) using 200 bins with a convergence tolerance of 1×10^-6^, and errors were estimated with 200 bootstraps. The WHAM method used considered the IACTs of each window, which were determined with the python module ‘pymbar’^22^. The value of each PMF was set to 0 in the region containing bulk water. The alternative PMF estimator, MBAR, was implemented using the python module ‘pymbar’^22^; PMFs calculated with MBAR used 100 bins. The free energy of lipid extraction was estimated by taking the difference between the minimum of the PMF and a point where the PMF had plateaued in solution.

### Simulation setup: Hamiltonian replica exchange

First, an OM was generated with the dimensions 8.6×8.6 nm^2^, and 4.0 nm above and below the membrane. The system was minimized and equilibrated for up to 30 ns and then the box height was increased to 16.8 nm. Once the system was re-solvated the minimization and equilibration steps were repeated. The equilibrium properties of the membrane were then converged over a 10 μs production run. All replicas were spaced using a geometric progression.

### Analysis

The ideal overlap between two harmonic biases spaced, *d*, apart was defined as,^21^

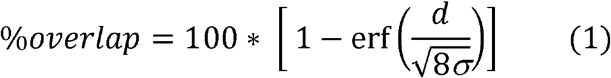

where *k* was the bias strength and *σ* was the gaussian strength, which was defined as:

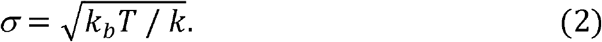

The tail order of each ReLPS lipid was determined using an in house script that was written with the python module ‘MDAnalysis’^48^. The average tail order of each acyl bond within 1 nm of the extracted lipid (in the xy plane) was then compared to that of a neat membrane. The deviations in order parameter were calculated for each bond in the acyl chain of ReLPS and defined as 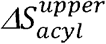. This analysis was based on previous work on the convergence equilibrium properties during the permeation of membranes by small molecules^13^. The orientation of the extracted ReLPS, θ_*LPS*_, was measured by taking the angles between the vector between the center of mass of the phosphate groups and the final sugar bead.

The APL and membrane thicknesses were determined with the software package FATSLiM^27^. The centers of geometry of the phosphate groups in each lipid were used as the reference points for the APL and membrane thickness calculations.

Some analysis was required to determine if LPS lipids were able to move past each other and thus properly mix. Two assumptions were made, the first one being that a single LPS should be neighbors with all other LPS lipids in a replica trajectory. The second assumption was that if LPS was properly mixing, then it should have an equal probability of being neighbors with any other LPS. To quantify both assumptions, nearest neighbor analysis was used and to track the six nearest neighbors of a single LPS across a trajectory. The first and second assumptions were quantified with the nearest neighbor index (NN index) and the *RMSD_NN_*, respectively. The NN index across a trajectory for a single lipid was defined as,

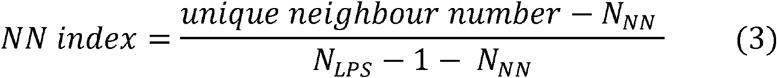

where *N_LPS_* was the number of LPS lipids and *N_NN_* the number of nearest neighbours in frame of the trajectory (i.e 6). If the NN index was 0 then the neighbours of an LPS lipid never changed. Conversely, if the NN index was 1 then the given LPS had been neighbours with every other LPS across the entire trajectory. The NN index referred to the results section was averaged across every LPS in the membrane.

The *RMSD_NN_* was defined for a single LPS as the RMSD of nearest neighbour frequencies, divided by the number of frames in a trajectory. If *RMSD_NN_* = 0 then the LPS had an equal probability of being neighbours with every other LPS across a trajectory. The *RMSD_NN_* referred to in the results section was averaged across all LPS lipids. The errors in the NN index and *RMSD_NN_* were determined from the standard deviation across all LPS lipids.

## Supporting information

all supplemental data

## Associated Content

### Supporting information

The supporting information is available free of charge on the ACS publication website at doi: xxx.

Contains further plots that were considered of minor importance to the main text (SUPP.pdf).

## Acknowledgments

For computational resources, we are grateful for continued support from the University of Southampton high performance computing resources, in particular for access to Iridis5, as well as the National Supercomputing Centre Singapore (NSCC) (https://www.nscc.sg). We are grateful to the UK Materials and Molecular Modelling Hub for access to Thomas, which is partially funded by EPSRC (EP/P020194/1) and HECBioSim for access to ARCHER through EPSRC (EP/R029407/1).

## Data availability

The data that support the findings of this study are available from the corresponding author upon reasonable request.

