## Supplementary material for "Molecular Dynamics Simulations of Bacterial Outer Membrane Lipid Extraction: Adequate Sampling?": all supplemental data

**Supplementary Methods**

**HREX: Trialling tempering regions**

For replica exchange, the number of replicas required for efficient sampling is proportional to the number of degrees of freedom in the tempered region^1^. Furthermore, the number of replicas for efficient sampling also scales with the size of the λ range. To minimize the computational cost of HREX MD, first a minimal tempering region must be identified. The trialed regions for which the forcefields were scaled are summarized in Figure S5. The largest tempering group was the entire ReLPS lipid (ALL). The rest of the tempered regions were comprised of parts of ReLPS: lipid A headgroup plus sugar moieties (HEADALL), the lipid A head group alone (HEAD), the sugar moieties alone (SUGARS), or the charged groups in LPS including all calcium ions in the system (CHARGED). Then 30 replicas, scaled by λ values geometrically spaced between 1 and 0.323, were simulated for 4 μs for each trial tempering region. No replicas were exchanged during these simulations, as the aim was to identify the ideal tempering region and range of λ values. Nearest neighbor analysis was used to determine that the ideal tempering region was the HEADALL region (Figure S6). The mixing when using the HEADALL region was only slightly less than using the entire lipid, even though the HEADLL group did not temper 38% of LPS beads. It is well known that the strength of the interactions between LPS lipids is highly dependent on the presence of divalent cations^2^, and it is noteworthy that tempering regions that consisted of only the charged beads yielded a smaller improvement in sampling than groups incorporating the headgroup sugars. The reason for this may lie in the well documented tendency for Martini carbohydrates to aggregate^3,4^.

**HREX: Covariance overlap**

The Block Covariance Overlap Analysis Method (BCOM) method can be used to quantify the relaxation speed of a system^5^. BCOM measures the covariance overlap between the PCA modes in different blocks of a trajectory. The sampling of a trajectory is sufficient when the PCA modes in each subset of the trajectory are the same and thus the covariance overlap is 1. The BCOM can be calculated as a function of increasing block size to ascertain whether the sampling is sufficient in a given trajectory. The subsets of a trajectory can also be generated through bootstrapping (BBCOM). The BCOM values can be normalized by dividing by the value of the frames in a trajectory that are uncorrelated i.e. the BBCOM value. The normalized BCOM as a function of increasing block size can be used to estimate the relaxation times of sampling in a system by fitting to a double exponential. Here the BCOM and BBCOM values were only calculated for the phosphates of ReLPS, as these were key positions that represent the lateral sorting of LPS and to reduce the computational expense of the analysis method. The BCOM analysis was applied across the last 10 μs of the ground state replica. For the sake of comparison, the unbiased ReLPS simulation was extended to 25 μs and the normalized BCOM determined across the last 10 μs (Figure S7). In both cases the block size was increased incrementally from 0.1 to 10 μs and 10 bootstrapped trajectory blocks were generated. The relaxation constants of the sampling in the HREX ground state were 0.13 and 1.25 μs, whereas the relaxation constants of the unbiased system were 0.20 and 1.55 μs. The normalized BCOM for the ground state was observed to smoothly approach 1, but did not reach 1 during the timescales sampled.

**HREX: Assessing efficiency of replica mixing**

The success of applying HREX MD to the OM was assessed though investigating efficiency of replica mixing. Here, replica mixing refers to the efficiency of the traversal of λ by all coordinate trajectories. In this context, a coordinate trajectory refers to a series of coordinates that are continuous in coordinate space, but not λ space; whereas a replica trajectory is continuous in λ space. The replica mixing was assessed across the entire 25 μs trajectory, and the exchange probability was determined for each replica (Figure S8A). The average exchange probability was ~0.36 and the range of the exchange probabilities was 0.07. The traversal of replica space was measured using the total residency time across replica space (Figure S8B). The replica mixing as a function of time was measured by calculating the transit fraction (Figure S8C). The transit fraction measured the fraction of coordinate trajectories that completed a minimum of 1 round trip as a function of time. The first round trip occurred after 4 μs, when the order of time for a round trip in previous atomistic simulations has been found to be on the order of nanoseconds^6–8^. By the end of the 25 μs only 40% of the replicas had experienced one round trip (Figure S8D). Using previous replica simulations^6^ as a benchmark, it is estimated that hundreds of microseconds of simulation would be required for sufficient mixing to be achieved.

**Supplementary Figures**

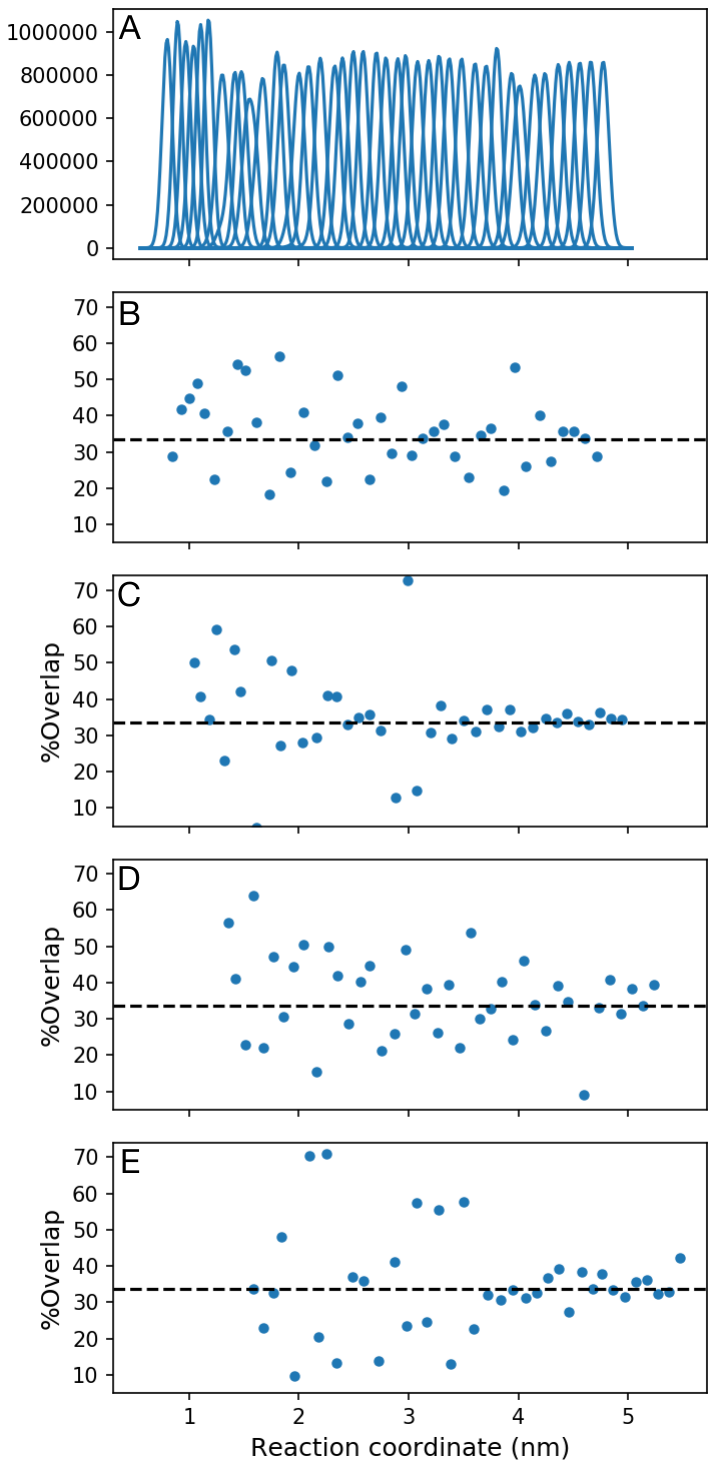

**Figure S1. US histogram overlap.** A) histogram of positions for 41 umbrella windows spaced every 0.1 nm for the COM_MEMB CV. Percentage overlap between adjacent windows for the extraction of ReLPS with the B) COM_MEMB, C) COM_CY D) HEAD_MEMB E) HEAD_CY CVs. The dotted line represents the ideal overlap between adjacent gaussians.

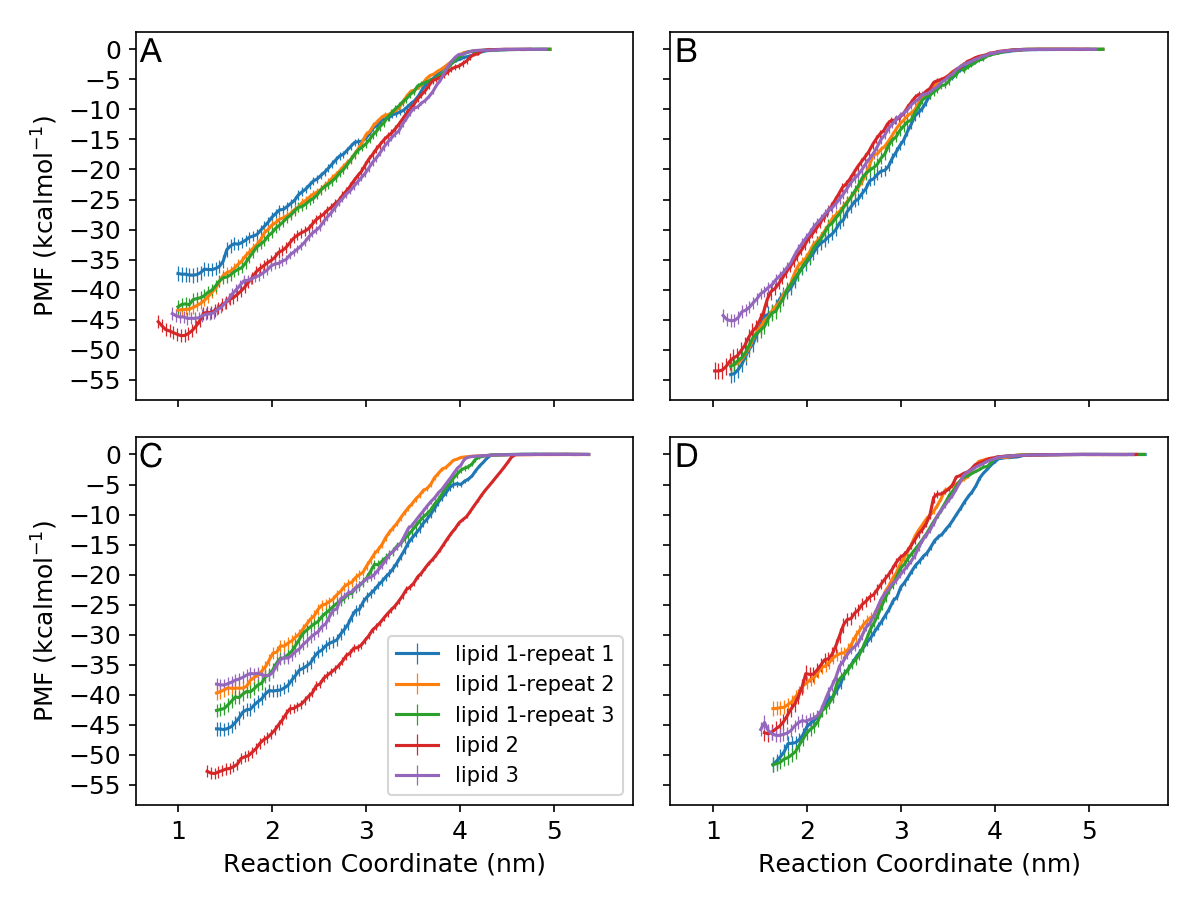

**Figure S2. PMFs for ReLPS extraction from the outer membrane.** PMFs are shown for A) COM_MEMB, B) COM_CY, C) HEAD_MEMB and D) HEAD_CY CVs, measured across the last 2 μs of each window. PMFs and errors were calculated using the MBAR free energy estimator.

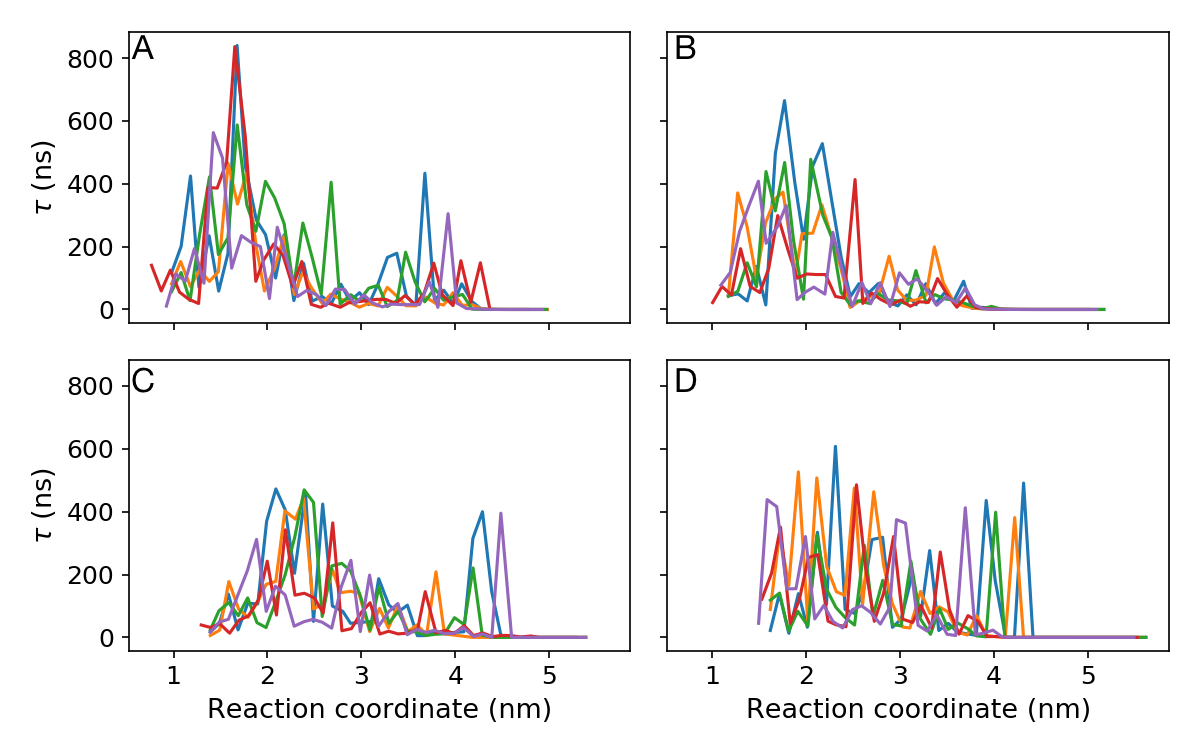

**Figure S3. Integrated autocorrelation times (τ) for each umbrella window of the extraction of ReLPS from the outer membrane.** Data are shown for A) COM_MEMB, B) COM_CY, C) HEAD_MEMB, and D) HEAD_CY CVs. Each line represents a different repeat.

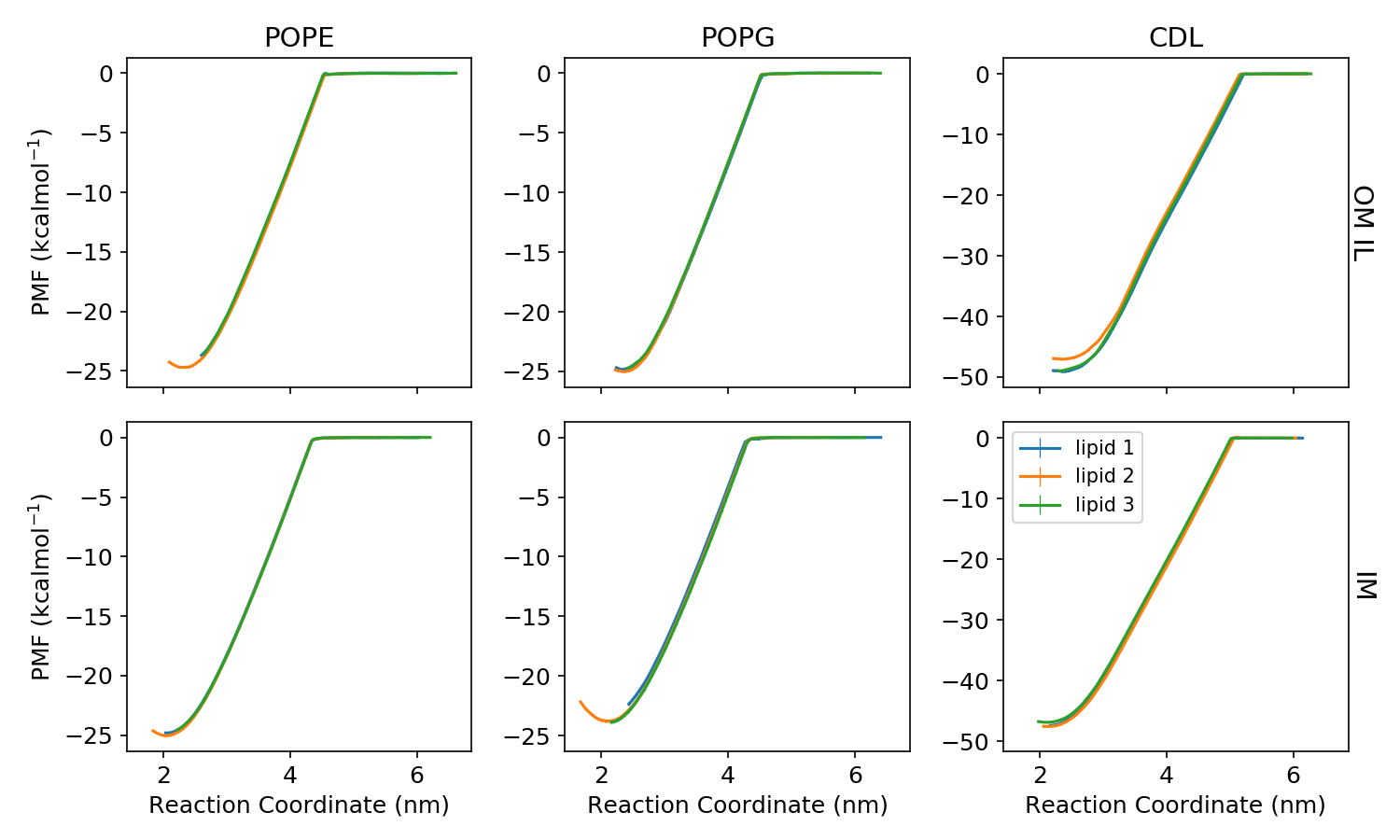

**Figure S4. PMFs for the extraction of POPE, POPG or cardiolipin (CDL) from either the OM inner leaflet (IL) or the IM.** PMFs were calculated over the last 0.5 μs of each umbrella window. Three different lipids were extracted per lipid type using the HEAD_CY CV.

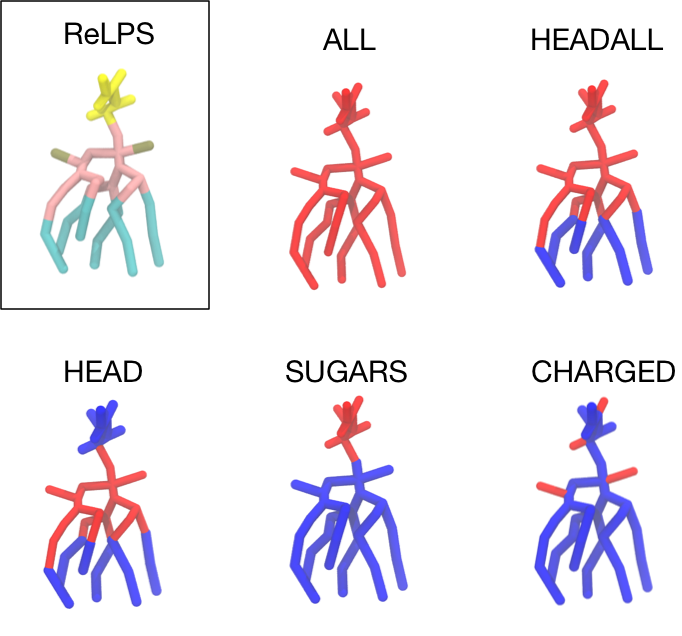

**Figure S5. HREX reference groups in LPS structures.** (Top left) CG structure of ReLPS (key: yellow = sugars, gold = phosphates, pink = lipid A sugar rings and acyl groups and cyan = lipid tails). The other structures depict the tempered regions (in red) trialed in this study. The tempered regions were as follows: ALL = entire lipid, HEADALL = sugars + lipid A headgroup, HEAD = lipid A headgroup, SUGARS = sugars, and CHARGED was any charged group on ReLPS and the calcium ions.

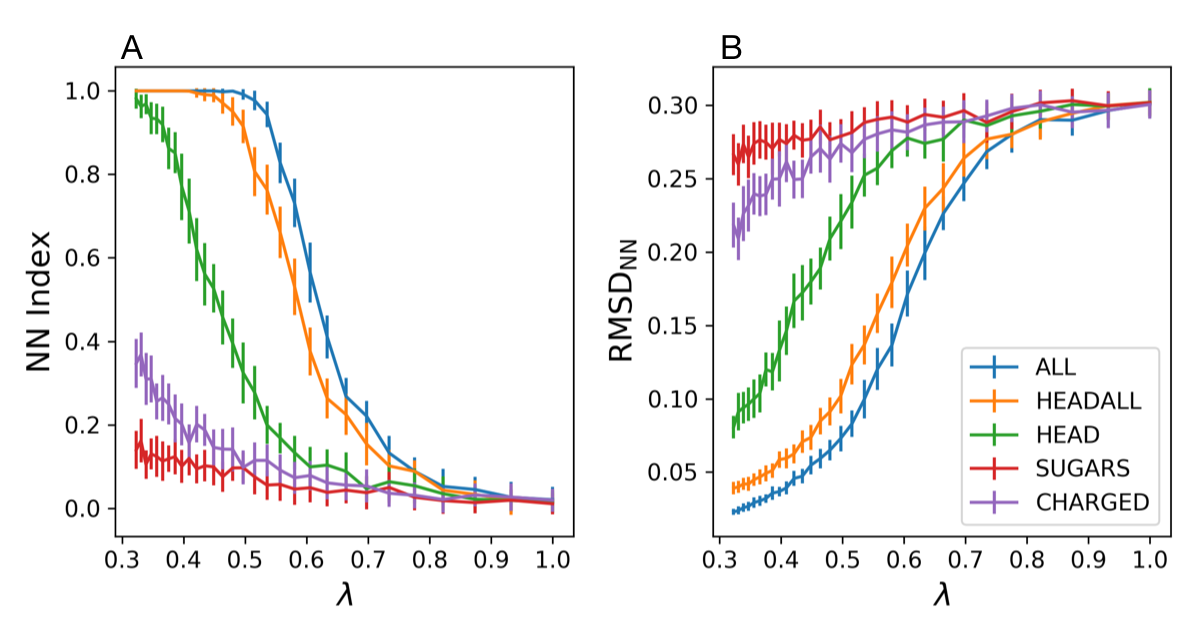

**Figure S6. Assessment of ideal tempering region during HREX MD simulations to enhance the lateral mixing of ReLPS in the OM model.** A) Nearest neighbor index (NN Index) and B) RMSD_NN_ vs λ for OM systems in which different regions were tempered. The regions that define each labelled tempering region can be found in Figure S5.

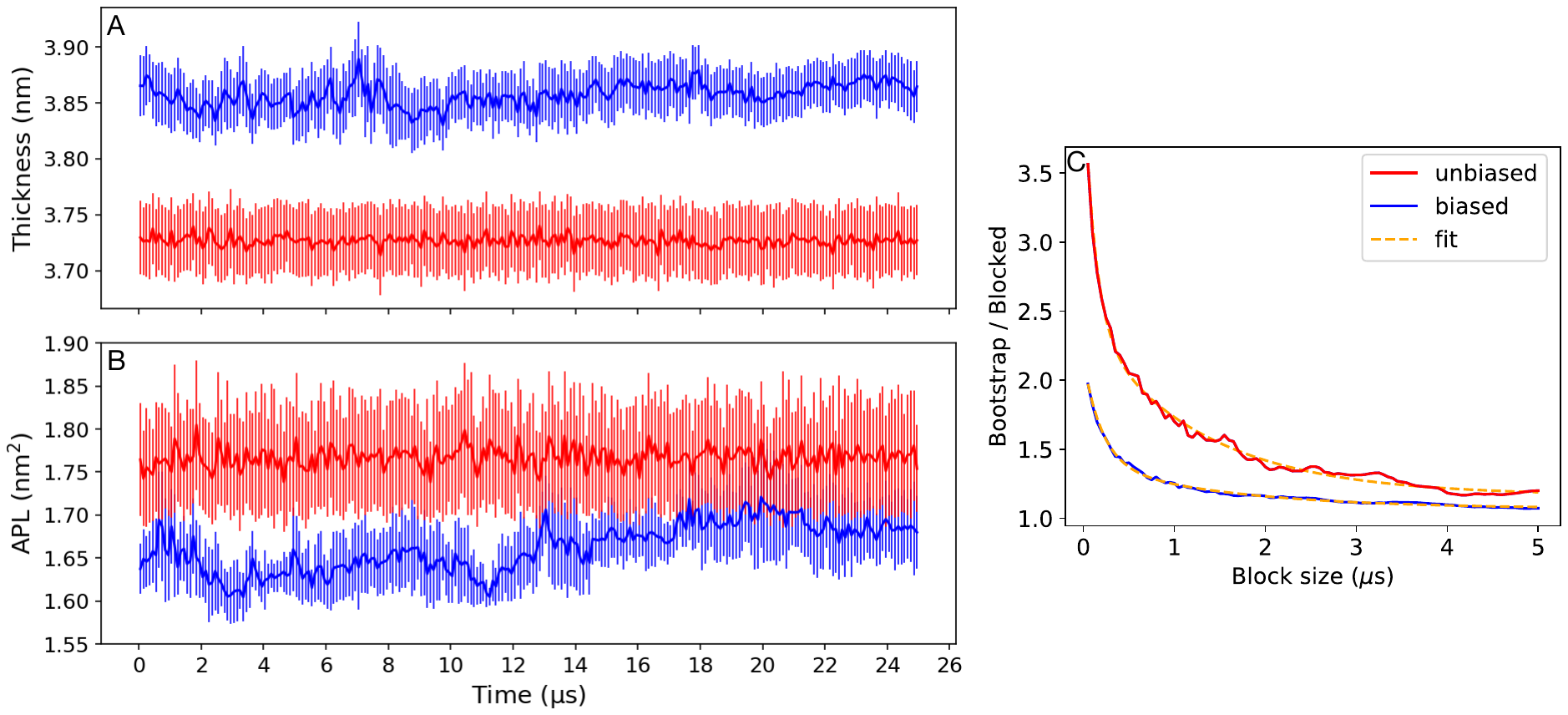

**Figure S7. Ratio between BCOM and BBCOM, based on ReLPS phosphate positions, during HREX simulations.**

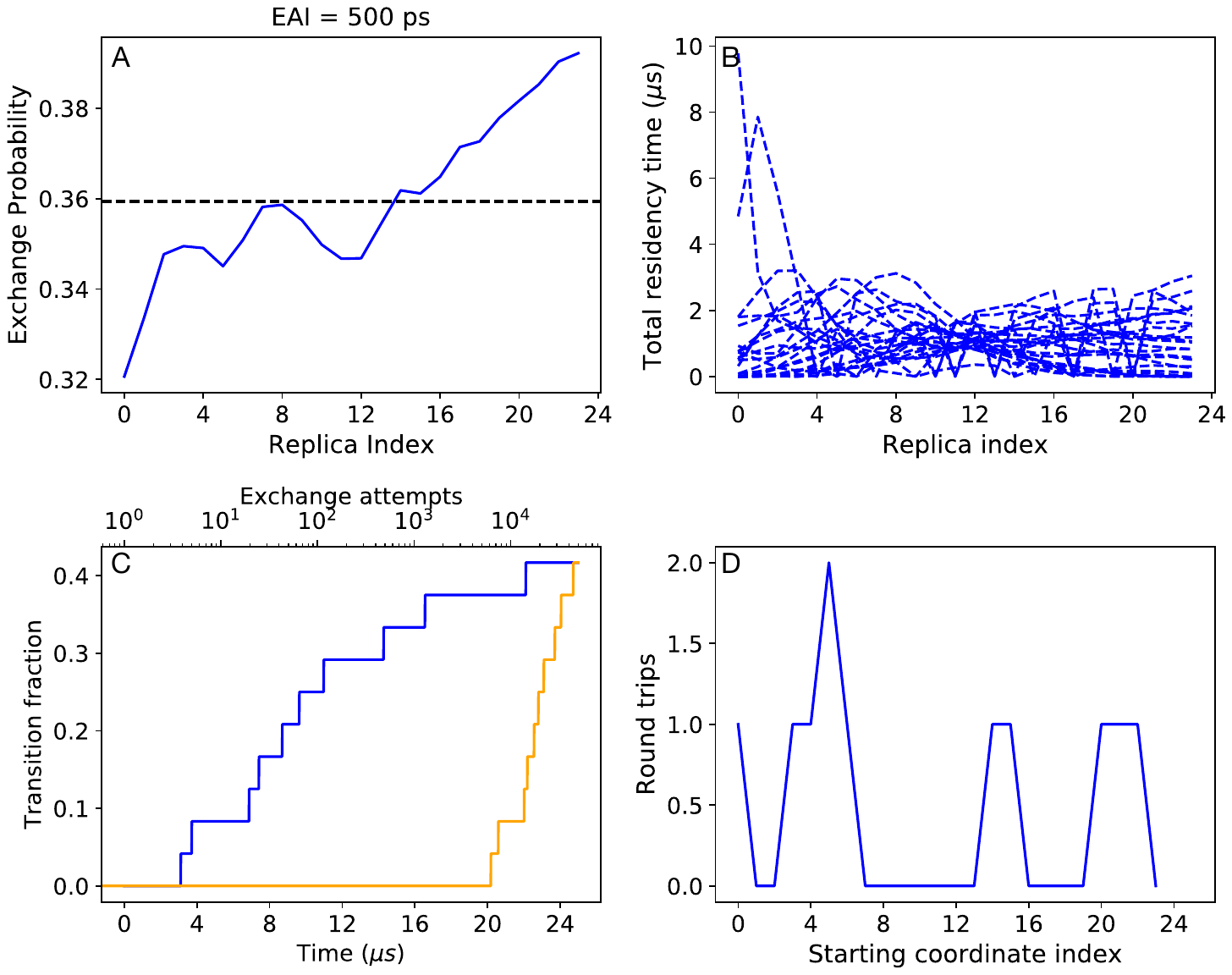

**Figure S8. HREX simulations, using the tempering of the HEADALL region in a ReLPS OM with exchange attempt intervals (EAI) of 500 ps.** A) The exchange probability of adjacent replicas as a function of the replica index. The dotted line indicates the average exchange probability. B) Total residency time for each coordinate trajectory (one coordinate trajectory per line) for every replica index in λ space. C) Fraction of replicas that complete at least one round trip vs time (blue) and number of exchange attempts (orange). D) Number of round trips for each coordinate trajectory vs starting coordinate index.
